# Lumped Parameter Simulations of Cervical Lymphatic Vessels: Dynamics of Murine Cerebrospinal Fluid Efflux from the Skull

**DOI:** 10.1101/2024.05.24.595806

**Authors:** Daehyun Kim, Jeffrey Tithof

## Abstract

**Background:** Growing evidence suggests that for rodents, a substantial fraction of cerebrospinal fluid (CSF) drains by crossing the cribriform plate into the nasopharengeal lymphatics, eventually reaching the cervical lymphatic vessels (CLVs). Disruption of this drainage pathway is associated with various neurological disorders.

**Methods:** We employ a lumped parameter method to numerically model CSF drainage across the cribriform plate to CLVs. Our model uses intracranial pressure as an inlet pressure and central venous blood pressure as an outlet pressure. The model incorporates initial lymphatic vessels (modeling those in the nasal region) that absorb the CSF and collecting lymphatic vessels (modeling CLVs) to transport the CSF against an adverse pressure gradient. To determine unknown parameters such as wall stiffness and valve properties, we utilize a Monte Carlo approach and validate our simulation against recent *in vivo* experimental measurements.

**Results:** Our parameter analysis reveals the physical characteristics of CLVs. Our results suggest that the stiffness of the vessel wall and the closing state of the valve are crucial for maintaining the vessel size and volume flow rate observed *in vivo*. We find that a decreased contraction amplitude and frequency leads to a reduction in volume flow rate, and we test the effects of varying the different pressures acting on the CLVs. Finally, we provide evidence that branching of initial lymphatic vessels may deviate from Murray’s law to reduce sensitivity to elevated intracranial pressure.

**Conclusions:** This is the first numerical study of CSF drainage through CLVs. Our comprehensive parameter analysis offers guidance for future numerical modeling of CLVs. This study also provides a foundation for understanding physiology of CSF drainage, helping guide future experimental studies aimed at identifying causal mechanisms of reduction in CLV transport and potential therapeutic approaches to enhance flow.

## Background

Cerebrospinal fluid (CSF) is a clear and colorless fluid that circulates around the brain and spinal cord, offering physical protection, acting as a “water cushion”to the central nervous system [1]. Additionally, it serves as a medium for supplying nutrients to the brain [2]. Beyond these traditional functions, CSF has gained recognition for its potential role in clearing metabolic wastes from the brain [3]. The discovery of meningeal lymphatic vessels [4], which permeate nearly the entire meninges (the layers of tissue surrounding the brain) in mammals, has significantly advanced research on CSF and its flow, potentially linked to various neurological disorders.

CSF is secreted from various sources, including the choroid plexus, ependymal cells, limited trans-capillary fluid flux, and metabolic water production [5]. Among these sites, the choroid plexus (found within the ventricles of the brain) is the main source, accounting for about 80% of CSF production. Following its production, CSF traverses the intricate ventricular system and subsequently reaches the subarachnoid space (SAS) surrounding the brain and spinal cord. Traditionally, the prevailing belief was that CSF is absorbed by arachnoid granulations. However, advancements in experimental techniques have revealed that substantial CSF also drains from the skull to eventually reach the cervical lymphatic vessels (CLVs) [6–11]. The contents of the CLVs may not be solely composed of CSF. The exact contribution of CSF to the total fluid within the CLVs is not fully understood and may vary under different conditions. However, it has been confirmed that fluid in the CLVs directly drains CSF, as microspheres injected into the cisterna magna of mice reach the CLVs within minutes [12].

The specific details of the route through which CSF reaches the CLVs are currently an area of active research and debate [13]. In animal studies, outflow along the cranial nerves, including the olfactory, optical, and facial nerves, as well as outflow to the meningeal lymphatic vessels and delivery to the deep cervical lymphatic nodes, has been well described. Compared to animal experiments, the understanding of CSF outflow pathways in humans is quite limited. Some evidence suggests that CSF outflow pathways exist through nasal lymphatics and meningeal lymphatic vessels [14, 15]. Several prior experiments in mice – especially of relevance for this study – suggests that a substantial fraction of fluid follows a pathway through the cribriform plate along the cranial nerves, draining into lymphatic vessels that eventually reach the CLVs [10, 12, 16–18].

Understanding fundamental aspects of CSF drainage to CLVs is significant due to its intricate association with various neurological disorders [19, 20]. Traumatic brain injury (TBI) also has been linked to CSF drainage disruption. In mouse models of TBI, research has revealed a significant reduction in CSF drainage through meningeal lymphatic vessels due to increased intracranial pressure (ICP) [21]. Additionally, it has been demonstrated that CSF flow serves as a conduit for transporting biomarkers associated with neurodegenerative diseases into the bloodstream via the cervical lymphatics [22]. More recently, TBI was linked to weakening of CLV activity and exacerbation of brain edema, which the authors argued arose as a consequence of a post-TBI surge in norepinephrine [12]. Notably, administration of a norepinephrine antagonist cocktail led to restoration of CLV function, reduction of edema, and improved cognitive outcomes. These findings suggest TBI may directly impact CLV function, disrupting CSF drainage from the skull.

Despite the significant implications for advancing disease treatments, precise quantification of CSF drainage routes has proven challenging. Spatial and temporal limitations in imaging pose one of the biggest challenges. Since lymphatic vessels are narrow and long, and traverse a substantial fraction of the body that can be difficult to optically access [23, 24], *in vivo* visualization of a large fraction of the lymphatic network that drains CSF is tremendously challenging. Moreover, the alteration of biological variables, such as ICP, during experimental testing has the potential to disrupt this delicate biological system [25]. As a consequence, these inherent constraints in experiments leave many open questions which can perhaps be addressed via numerical simulation.

Numerical simulations serve as efficient, safe, and ethically desirable alternatives to experiments; simulations also enable manipulation of parameters and the ability to prescribe/determine precise conditions that may be difficult or impossible to achieve experimentally, such as pressures inside vessels. Numerous prior studies have used a numerical approach to model lymphatic vessels. Reddy et al. pioneered this field by creating the initial mathematical model for branching lymphatic vessels [26]. Their one-dimensional (1D) model simulated laminar lymphatic flow throughout the entire body, from the periphery through the main lymphatic system into the venous blood system. Their governing equation was based on the Navier-Stokes equation (in a 1D form) coupled with a thin-wall tube model. Quick et al. employed a circuit-theory approach, commonly employed in the simulation of cardiovascular systems [27], to simulate both individual lymphangions and a chain of lymphangions. Bertram et al. extended this methodology and focused on a series of lymphangions, highlighting the efficiency of sequential contractions compared to synchronized contractions for fluid transport [28]. These foundational models lay the groundwork for simulating a large number of interconnected lymphangions, with incorporation of compliant walls and valves. Notably, these models were primarily based on parameters from the thoracic duct and mesenteric lymphatic vessels.

In parallel, two-dimensional (2D) or three-dimensional (3D) models have typically focused on a single lymphangion and often utilize fluid-structure interaction techniques [29–31]. Such studies have investigated the influence of active contractions driven by nitric oxide and calcium ions on fluid drainage. Validation of Poiseuille flow under substantial radial contractions was also studied. Computational cost has largely confined such studies to simulation of a single lymphangion, although some 2D/3D simulations for a series of lymphatic vessels, leveraging fluid-structure interaction approaches, have been achieved [32– 34]. These studies tackled a range of factors, including the chain’s length and its influence on mean flow rates, the effects of adverse pressure differences on flow rates, contraction timing, dynamics related to valve geometry impacting backflow, and the effects of valve elasticity. Again, it is worth highlighting that these simulations were based on lymphatic vessels in the other regions, such as mesenteric lymphatic vessels, and were not specifically tailored to CSF drainage through CLVs.

Currently, there is an absence of numerical models that predict fluid transport through CLVs. The goal of this study is to construct a comprehensive lumped parameter model that captures the intricate CSF drainage process in mice. In the absence of knowledge of the exact geometry and physical properties of CLVs, we use a Monte Carlo approach to identify a parameter regime that matches closely with experimental data. Simultaneously, we perform a parameter sensitivity analysis to examine the characteristics of CLVs and how changes in these parameters affect the CLV diameter and net volume flow rate. Using our model, we explore strategies to enhance CSF flow rate through CLVs by manipulating certain variables. Finally, we explore how the variation of ICP affects CSF efflux through the CLVs. This study is the first numerical model of CSF outflow through CLVs, providing novel insights into potential strategies to enhance CSF drainage. These findings provide guidance for future experimental research and serve as a valuable resource for optimizing CSF drainage strategies.

## Methods

We use the lumped parameter method to model the nasal and cervical lymphatic system, which are treated as initial and collecting lymphatics. The versatility and computational efficiency makes this approach suitable for simulating the complexities and capturing the impact of uncertainties associated with this lymphatic network that connects the SAS to the end of the CLVs. Given the lack of detailed parameter values for CLVs in the literature (e.g., stiffness of the vessel wall, active tension due to smooth muscle cells, valve properties), this approach allows for computationally-efficient iterative determination of parameters to obtain agreement with experimental observations.

### Modeling CSF Drainage via CLVs

Lymphatic vessels typically consist of initial lymphatic vessels (lymphatic capillaries) and collecting lymphatic vessels [35]; in this study, CLVs are considered the only collecting lymphatic vessels. The initial lymphatic vessels form the starting point of the lymphatic system and are responsible for absorbing ISF. These vessels consist of a single layer of endothelial cells, and their button-like junctions structure allows ISF and small particles to enter. As these interstitial lymphatic vessels merge, they give rise to larger lymphatic vessels known as collecting lymphatic vessels. Collecting lymphatic vessels consist of interconnected compartments called lymphangions which serve as the primary transporters of lymph within the lymphatic system. Unlike initial lymphatic vessels, collecting lymphatic vessels are covered by smooth muscle cells. There are two types of pumping mechanism in the collecting lymphatic vessels: extrinsic and intrinsic. Extrinsic pumping primarily occurs due to external forces acting on the lymphatic system. These external forces include skeletal muscle contractions, external compression from the pulsation of nearby arteries, and/or movement caused by respiration. Intrinsic pumping refers to the transport arising from natural contractile activity of smooth muscle cells lining the basement membrane of lymphatic vessels. In the periphery, a combination of extrinsic and intrinsic forces facilitate the movement of lymph against an adverse pressure gradient. The presence of valves between lymphangions ensures net transport of lymph by preventing backflow. Thus intrinsic and extrinsic pumping enables collecting lymphatic vessels to efficiently transport lymph to lymph nodes.

We used this general lymphatic structure to model CSF outflow through CLVs. A schematic of the model is illustrated in Fig. 1A. Due to limited anatomical data and flow measurements for humans, we instead developed our model to capture murine anatomy. Even though the exact route connecting the CSF in SAS to the lymphatic capillaries is not fully established [13], recent experiments suggest that tiny lymphatics have direct connection to the SAS across the cribriform plate [16, 36–38]. Based on these findings, we assume there are direct connections through which the lymphatic capillaries absorb CSF from the SAS. These capillaries then merge to form larger vessels, with diameters prescribed by Murray’s Law but with a different exponent value (*n* = 1.45) [39]. For simplicity, we assume that initial lymphatics are impermeable and absorption of CSF occurs at the tip of the vessels. CSF in the initial lymphatics is transported to the CLVs, then eventually pumped to the central venous blood. Fig. 1B summarizes the mathematical model that describes CSF efflux from the SAS to CLVs then venous blood. Our model is based on the lumped parameter method, which is analogous to an electrical circuit, so our simulation diagram includes circuit elements, including an equivalent resistor *R*_*initial*_ which captures the hydraulic resistance of the initial lymphatic vessels.

**Fig. 1:**
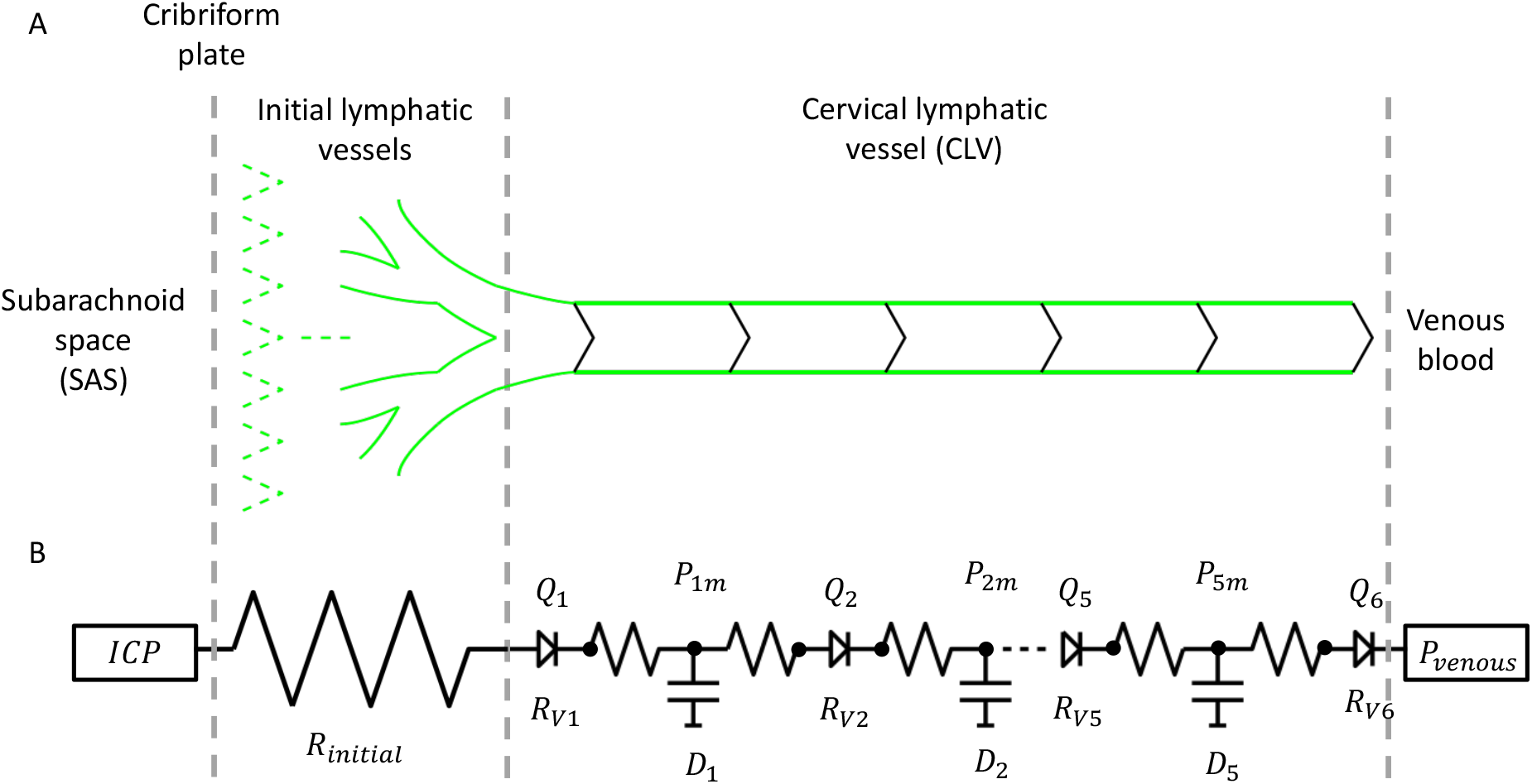
Comparative illustration and mathematical approach to modeling CSF drainage. (A) Illustration of the biological pathway for CSF drainage through CLVs. CSF in the SAS is directly absorbed by lymphatic capillaries that cross the cribriform plate. These capillaries merge to form larger vessels, which eventually connect to the CLVs. The CLVs then transport the CSF back into the central venous blood. (B) Schematic of the electrical model of this CSF drainage pathway. The inlet pressure is the ICP, and the lymphatic capillaries are modeled using a net equivalent hydraulic resistance (*R*_*initial*_). The outlet pressure is equal to that of central venous blood. CLVs are modeled to incorporate contractions and expansions of lymphangions, as well as opening and closing of valves in between. Hydraulic resistance of the valves (*R*_*V*_) are represented as LEDs in the electrical circuit. Each lymphangion has the same length (*L*) and its own diameter (*D*_*j*_). While capacitance is not explicitly included in our governing equations, the change in vessel diameter serves a capacitance-like function and is therefore denoted with a capacitance symbol. The pressures within each lymphangion are calculated at three points: just past the inlet, the midpoint, and just before the outlet. The inlet and outlet pressures are used to calculate the volume flow rates (*Q*) at the valves. The pressures at the midpoint (*P*_*m*_) are determined using the transmural pressure equation. The design of the CLV part is inspired by Bertram et al. [28].

The size of the initial lymphatics located near the cribriform plate was estimated by analyzing images from Spera et al. [18] (tip diameter: 7.5 µm) and Norwood et al. [16] (length of total branches: 370 µm). By applying the modified Murray’s Law 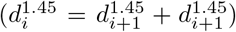, we calculated the diameters of parent vessels and number of branching generations until their diameter reached the median diameter of CLVs (*D* = 84.1 µm) as reported in Hussain et al. [12]. We assume that the tip of two daughter branches of the lymphatic capillaries with the same diameter merge into one parent vessel. Eventually, the merged parent vessels join to form the first lymphangion of the collecting lymphatic vessels. We obtained *N* = 5 for the number of generations (compared to *N* = 11 for standard Murray’s law with exponent 3). The modified exponent in our model results in a smaller generation (*N* = 5), leading to smaller diameters of the branches. Also, each branch becomes longer, as the total length of the entire branching structure is fixed. Subsequently, we computed the hydraulic resistance of this branching network under the assumption of Hagen-Poiseuille flow. Finally, the total hydraulic resistance was calculated using computation of series of parallel resistances as,

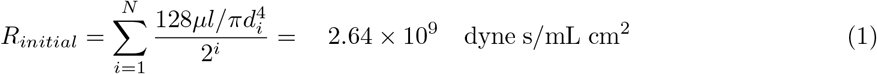

where *i* is the specific generation, *µ* is dynamic viscosity of the CSF, *l* is length of each segment of daughter capillaries, and *d*_*i*_ is the initial lymphatic vessel diameter at generation *i*. Note that lower case *d* and *l* are used to represent the initial lymphatic vessels, while upper case *D* and *L* are used to represent the collecting lymphatic vessels.

Bertram et al’s works are the inspiration for our approach to modeling CLVs [28, 40, 41]. We assume a collecting lymphatic vessel consists of five lymphangions, based on our estimates of 10 mm from the nasal mucosa to superficial cervical lymph node and a 2 mm lymphangion length [12, 17]. Each lymphangion, *j*, is characterized by time-dependent variables, namely, CLV diameter *D*_*j*_, and pressures *P*_*j*,1_, *P*_*j,m*_, and *P*_*j*,2_ representing the pressures just after the inlet, at the midpoint, and just before the outlet, respectively. Note that the use of three points per lymphangion (rather than just one) enables more realistic modeling that can capture valve prolapse [42]. Additionally, the model includes the volume flow rates *Q*_*j*_ and *Q*_*j*+1_, representing the inflow and outflow rates at each valve of a given lymphangion, respectively. The fluid flow from one chamber to the next chamber is described by a control volume discretization of the equations governing conservation of mass and momentum [28, 40, 42],

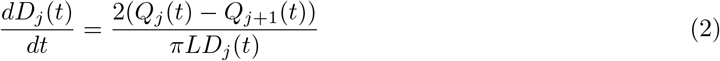

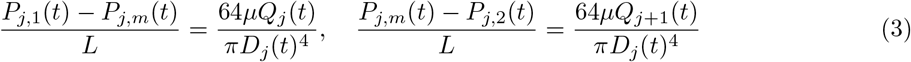

under the quasi-steady assumption of fully developed Hagen-Poiseuille flow. This assumption is justified by the small Reynolds number (*Re* = *UD*_0_*/ν* ≈ 7 *×* 10^−3^) and small Womersley number 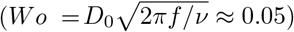 of flow in the CLVs, where *U* = 64.8 µm/s is the mean flow speed, *D*_0_ = 84.1 µm is the mean CLV diameter, *ν* = 0.7 × 10^−6^ m^2^/s, and *f* ≈ 0.04 Hz is the contraction frequency. Note that we provide derivations of equations (2-3) in Appendix A. Each lymphangion is separated by valves with hydraulic resistance [28],

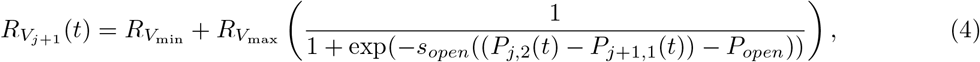

where 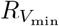 and 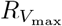 are the respective minimum and maximum hydraulic resistance of the valves when open or closed, *P*_*open*_ is the minimum pressure required to open the valve, and *s*_*open*_ is the slope of the valve opening. A large *s*_*open*_ means a more rapid response of the valve opening to changes in pressure difference across the valve, while a smaller *s*_*open*_ indicates a more gradual response. The volume flow rate through the valves is calculated based on a momentum equation analogous to Ohm’s law,

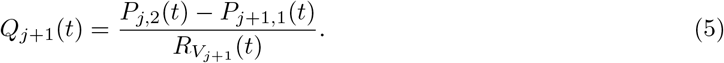

The associated sets of differential algebraic equations are then closed by specifying the transmural pressure (i.e., pressure difference between inside and outside the lymphangion) using an equation that contains both passive and periodic contractile components [28]:

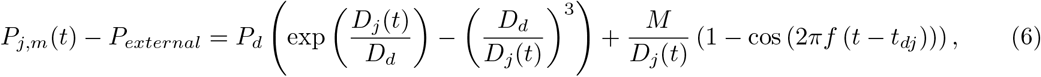

where *P*_*external*_ is a uniform pressure external to the lymphangions, *P*_*d*_ and *D*_*d*_ are passive properties that account for mechanical characteristics of lymphangion’s contractions and expansions, *M* is the active tension, and *t*_*d*_ is temporal phase between contractions of adjacent lymphangions (these parameters are discussed in more detail below). We assume the external pressure *P*_*external*_ is constant and slightly lower than the inlet and outlet pressures to make the transmural pressure operate more dynamically, similar to Bertram et al’s work [28].

### Parameter Estimation

We first discuss parameters in the equations that are well-known based on literature and experimental observations, which are detailed in Table 1. The frequency of lymphangions’ contractions, *f*, and the length of each lymphangion, *L*, are determined directly from in vivo images [12] using our recently published image analysis techniques [43]. The dynamic viscosity of CSF was used in our model [44]. We used the same values for the temporal phase between adjacent lymphangion contractions, *t*_*d*_, and the slope of valve opening, *s*_*open*_, as in previous studies [28, 41].

**Table 1:** Reasonably well-known parameters for modeling CLVs, as used in this study.

| Parameters | Definitions | Value | Unit | Reference |
| --- | --- | --- | --- | --- |
| $f$ | Contraction frequency | 2.4<br>(Default)<br>1 - 29<br>(Fig. 5B) | $\text{min}^{-1}$ | [12] |
| $L$ | Length of lymphangion | 2 | mm | [12] |
| $n$ | Number of lymphangions | 5 | | Assumed |
| $t_d$ | Temporal phase between adjacent lymphangion contractions | 90 | degree | [41] |
| $s_{open}$ | Slope of valve opening | 0.2 | $\text{cm}^2/\text{dyne}$ | [40] |
| $\mu$ | Dynamic viscosity of CSF | 0.007 | $\text{Pa} \cdot \text{s}$ | [44] |
| $P_{external}$ | External pressure | 3.75<br>(Default)<br>2 - 4<br>(Fig. 5C) | mmHg | Assumed |
| $ICP$ | Intracranial pressure | 4<br>(Default)<br>10<br>(Elevated ICP for Fig. 6)<br>2.5 - 7<br>(Fig. 5D) | mmHg | [46, 47] |
| $P_{venous}$ | Central venous pressure | 6<br>(Default)<br>4.16 - 8.94<br>(Fig. 5D) | mmHg | [48] |
| $R_{initial}$ | Equivalent hydraulic resistance of lymphatic capillaries | $3.21 \times 10^8$<br>(n = 3)<br>$2.653 \times 10^9$<br>(n = 1.45) | $\text{dyne} \cdot \text{s}/(\text{mL} \cdot \text{cm}^2)$ | [39] |

Estimating other parameters poses a substantial challenge, particularly for those not directly measurable, such as properties of the cervical lymphatic vessel walls (*P*_*d*_, *D*_*d*_), active tension generated by smooth muscle cells (*M*), the minimum pressure required to open the valve (*P*_*open*_), and hydraulic resistance of the valves 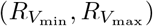. While more detailed explanation of these constants is available in the original works [28, 40, 41, 45], it is worthwhile to provide a brief explanation of these unknown properties here. *P*_*d*_ represents the vessel wall stiffness. A higher *P*_*d*_ value means that the transmural pressure reacts more sharply to any change in vessel diameter. *D*_*d*_ is the threshold diameter at which the vessel begins to experience positive or negative transmural pressure. When the lymphangion’s diameter (*D*_*j*_) surpasses *D*_*d*_, the transmural pressure becomes positive, with the shape of an exponential curve. Conversely, when the lymphangion’s diameter is less than *D*_*d*_, the transmural pressure becomes negative, described by a cubic function. *M* represents the tension generated by smooth muscle cells around the vessel. The transmural pressure incorporates this active tension (*M*) as well as the vessel’s passive properties (*P*_*d*_ and *D*_*d*_) to account for the mechanical characteristics of lymphangion’s contractions and expansions. *P*_*open*_ is the threshold pressure for the opening of valves between adjacent lymphangion. The valves are analogous to a diode in an electrical circuit. When the pressure difference across the valve exceeds *P*_*open*_, the resistance of the valve becomes approximately 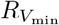. When the upstream pressure (pressure before the valve) is lower than the downstream pressure (pressure after the valve), the hydraulic resistance of the valve becomes approximately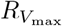, meaning that if this value is very large, the valve is practically closed.

To overcome the challenge of not knowing the mechanical properties of CLVs, we adopt an inverse problem-solving approach based on a Monte Carlo method in which we vary parameters then compare model outputs (diameter of the third lymphangion and volume flow rate through the downstream valve of the third lymphangion) to experimental measurements of the median vessel diameter (84.072 µm) and mean volume flow rate (0.0226 µl/min) [12]. This approach allows us to iteratively adjust and optimize our parameters by validating against the known outputs, aiming to achieve a more accurate and comprehensive model of the CSF drainage dynamics.

We set bounds for unknown parameters based on prior research on rat mesenteric lymphatic vessels [28, 40, 41, 45, 49]. The lower and upper bounds we set come from the minimum and maximum of reported values. We then expanded the bounds by subtracting or adding twice the standard deviation *σ*. Our resulting bounds for unknown parameters are summarized in Table 2. We next generated a random uniform distribution for each of the unknown parameters in accordance with each range. 10,000 simulations were conducted based on these randomly sampled parameters, and results from the model were compared to the median diameter and mean volume flow rate of the CLVs as measured *in vivo*.

**Table 2:** Bounds on unknown parameters, used in our Monte Carlo approach to modeling CLVs to determine parameter sets that lead to simulations that closely match average experimental measurements.

| Parameters | Definitions | Lower Bound | Upper Bound | Unit |
| --- | --- | --- | --- | --- |
| $M$ | Active tension | 3.6 | 24.3 | dyne/cm |
| $R_{V_{min}}$ | Minimum valve resistance | 600 | $2.83 \times 10^6$ | dyne · s/(mL · cm<sup>2</sup>) |
| $R_{V_{max}}$ | Maximum valve resistance | $1.2 \times 10^7$ | $2.82 \times 10^{10}$ | dyne · s/(mL · cm<sup>2</sup>) |
| $p_{open}$ | Valve opening pressure | -70 | -15 | dyne/cm<sup>2</sup> |
| $P_d$ | Constitutive-relation constant | 20 | 732 | dyne/cm<sup>2</sup> |
| $D_d$ | Constitutive-relation constant | 0.00845 | 0.0613 | cm |

Diameters, pressures, and volume flow rates in the differential algebraic equations (1)-(3) coupled with valve and transmural pressure equations (4)-(6) were solved numerically. ‘*fsolve*’ function from the SciPy library was used to find the transmural pressures of each lymphangion. Diameters were calculated by employing the 4th order Runge-Kutta method. These calculations continued until converged periodicity of variables was achieved. Finally, the volume flow rates at the valves were calculated based on the hydraulic resistance of the valve and pressure difference across it. We computed the median diameter *med*(*D*_*sim*_) and mean volume flow rate 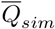 for each simulation, which we used as criteria for the search. If the comparison fell within 5% of the experimental value (denoted with subscript “exp”), such that,

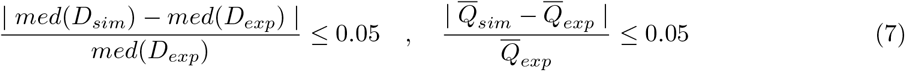

we noted that particular parameter set. We continued this process for every randomly chosen parameter set and repeated the same numerical procedure. A summary of this process, including the Monte Carlo parameter selection, is depicted in Fig. 2. Note that we stored the results of all simulations (including those that did not meet our 5% criteria) for the sake of performing sensitivity analysis.

**Fig. 2:**
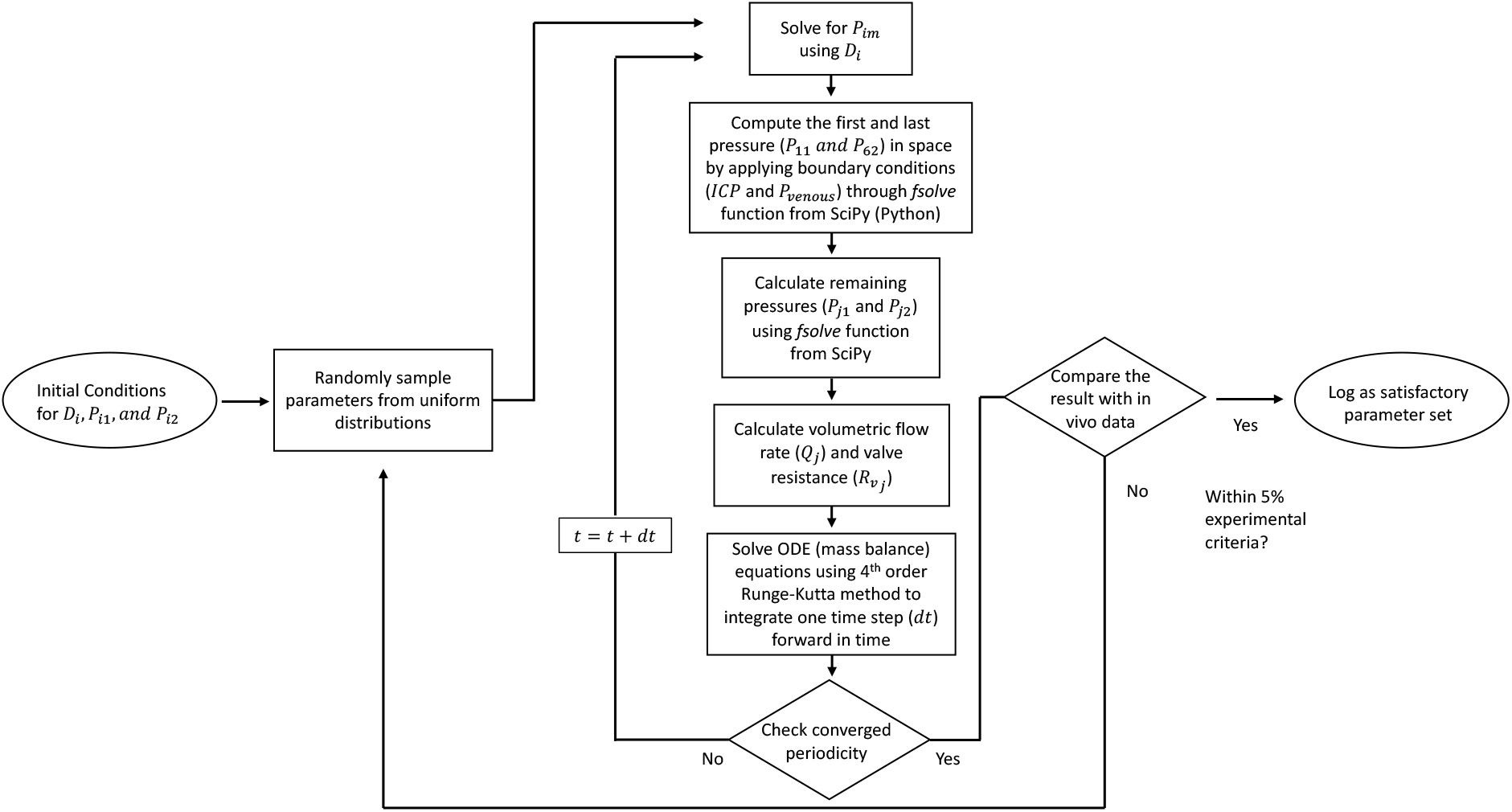
A flow chart depicting the algorithm used in our Monte Carlo parameter search.

## Results

We now present simulation results, which detail our parameter estimations, overall CSF efflux through lymphatic vessels, a sensitivity analysis of the parameters affecting CLVs, and the impact of branching in initial lymphatic vessels. It is important to note that the volume flow rate is calculated at the valves, whereas the diameters are calculated in the middle of each lymphangion. The mean volume flow rate 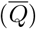 used throughout the results refers to the time-averaged downstream volume flow rate of the third lymphangion in our simulation. Similarly, the reported median diameter is the temporal median of the third lymphangion diameter. Additionally, we restate the definition of each unknown parameter used in the Monte Carlo simulation, for convenience. *P*_*d*_ and *D*_*d*_ represent the properties of the CLV wall that control the stiffness of the vessel. *M* denotes the magnitude of active tension generated by smooth muscle cells. *P*_*open*_ is the minimum pressure required to open the valve. 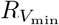 is the hydraulic resistance of the valves in their open state, whereas 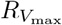 is the hydraulic resistance of the valves in their closed state.

### Inverse Problem: Monte Carlo Approach

We conducted a Monte Carlo simulation to estimate parameters that are consistent with experimental observations. Out of 10,000 simulations, only 585 successfully converged. The converged simulations indicate that *M, D*_*d*_, and *P*_*d*_ must fall within a narrow range for simulations to successfully converge. To further explore the sensitivity, we narrowed the values of *M, D*_*d*_, and *P*_*d*_ and performed an additional 2,000 simulations, randomly sampling parameters from within these revised limits (Fig. 3).

**Fig. 3:**
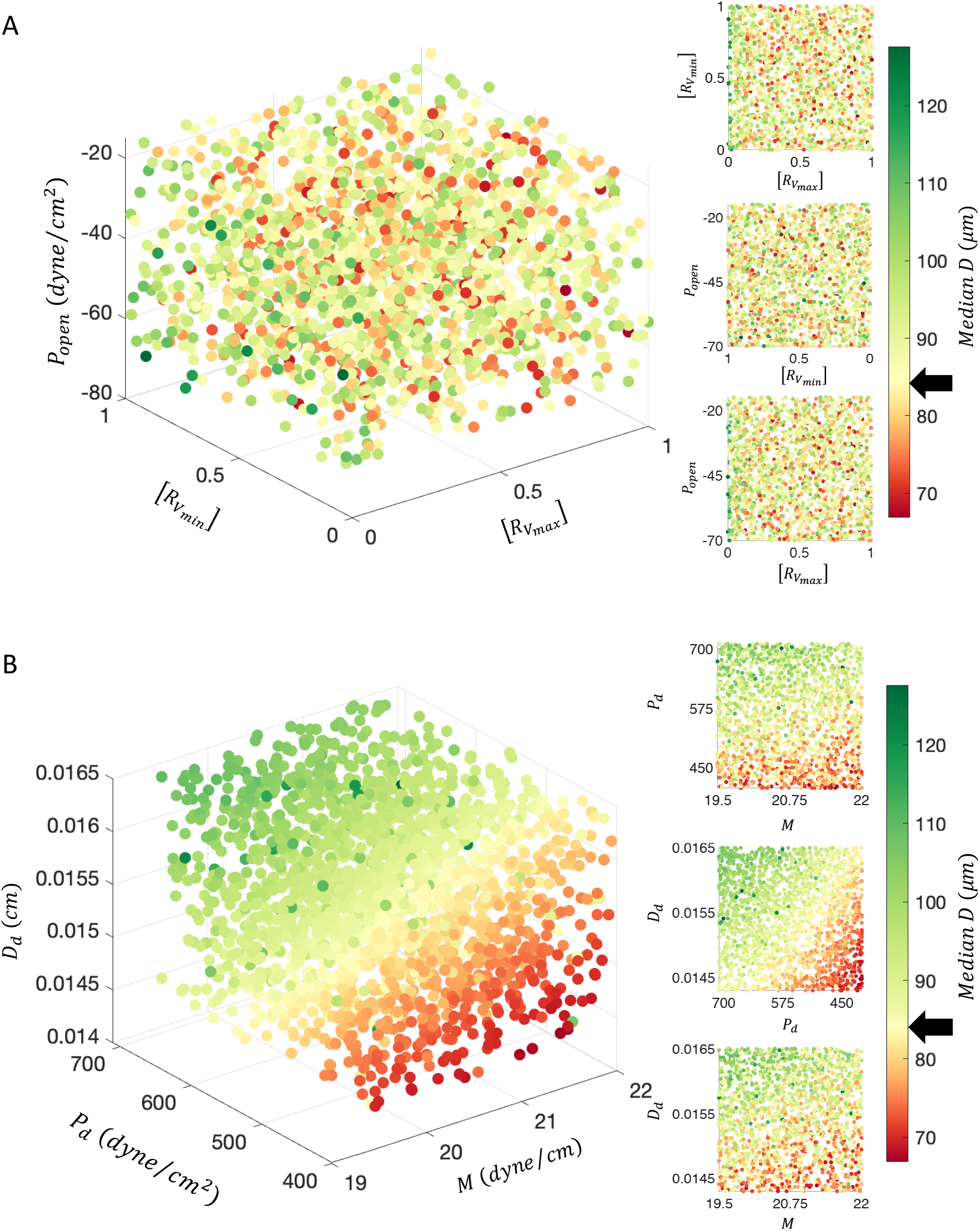
Results from the Monte Carlo search and analysis of parameter sensitivity (figure part one of two). The large 3D scatter plots on the left side illustrate the overall sensitivity to the chosen parameters on each axis. The smaller three plots on the right provide 2D projections onto each pair of axes. The target values for each criteria are indicated by black arrows on the color bars. (A) 3D scatter plots showing the variation of median diameter with changes in 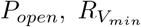, and 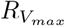. The hydraulic resistance of the valves, 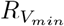 and 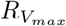 are normalized by their maximum values such that 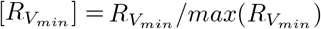 and 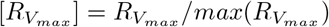(see text for values). (B) 3D scatter plots showing the variation of median diameter with changes in *D*_*d*_, *P*_*d*_, and *M*.

**Fig. 3:**
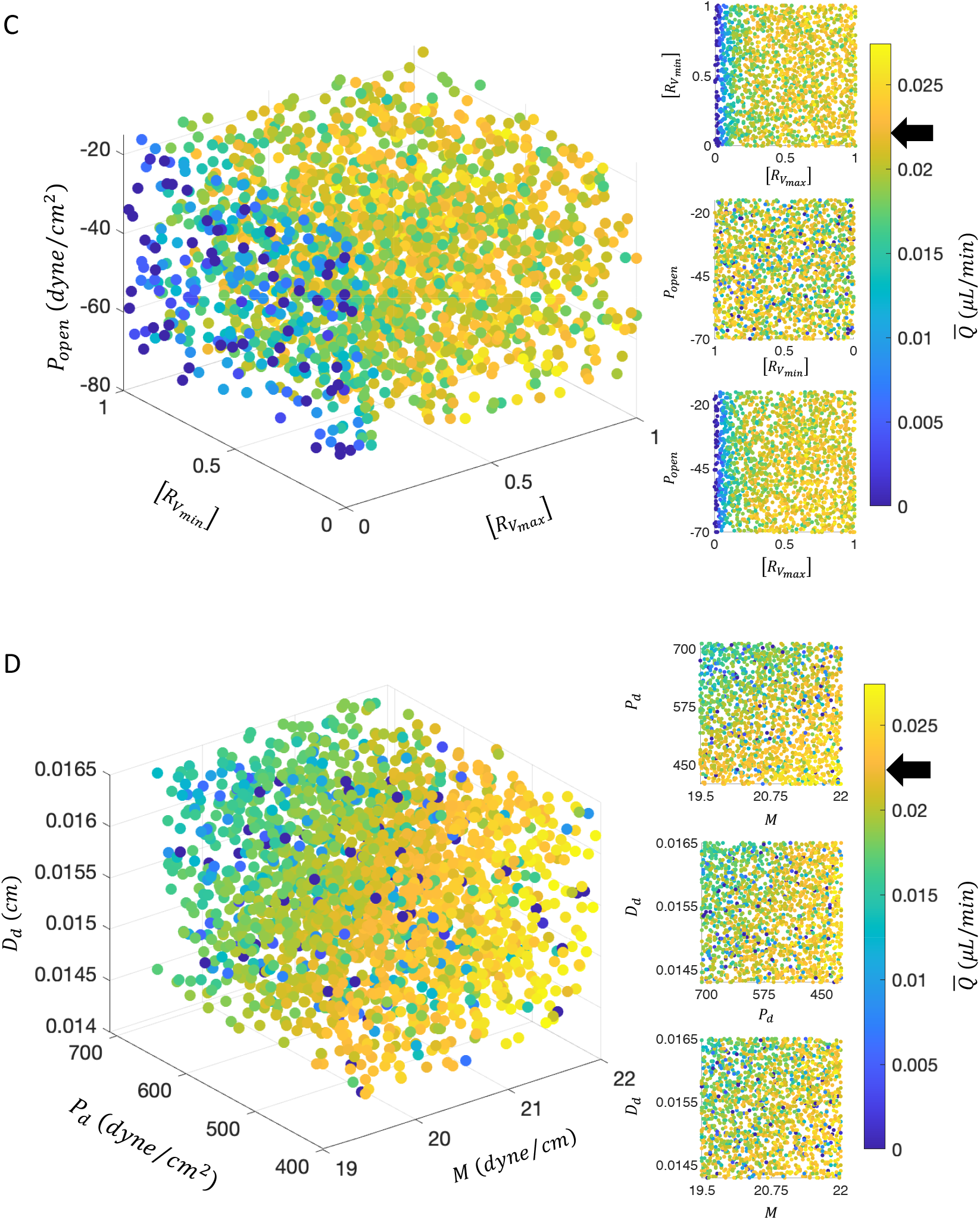
Results from the Monte Carlo search and analysis of parameter sensitivity (figure part two of two). (C) 3D scatter plot showing the variation of mean volume flow rate 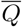 with changes in 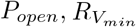, and 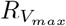. (D) 3D scatter plot showing the variation of mean volume flow rate with changes in *D*_*d*_, *P*_*d*_, and *M*. For (C) and (D), volume flow rates below zero are plotted as zero to emphasize the sensitivity among positive values.

The parameter sensitivity analysis revealed that *P*_*open*_ and 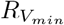 do not significantly impact the vessel diameter, as no clear trend in median diameter value emerged as each of these parameters were varied (Fig. 3A and Supplementary Fig. S1B, D). Values close to the experimental value we sought to match (84.1 µm), in a color indicated by the black arrow on the color bar, are randomly distributed throughout the parameter space. 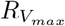 had a barely discernible influence on the diameter when 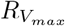 was very small; such values led to increases in diameter of around 120 µm (see green circles at left in Fig. 3A). On the other hand, the parameters *D*_*d*_, *P*_*d*_, and *M* were found to influence vessel size, with a strong tendency for the diameter to increase with lower *M*, higher *P*_*d*_, and higher *D*_*d*_ (Fig. 3B and Supplementary Fig. S1A, E, F). This indicates that the balance between the stiffness of the vessel wall and the active tension exerted by smooth muscle cells significantly affects the diameter of the CLV.

The mean volume flow rate was not greatly affected by *P*_*open*_ or 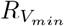, but showed significant sensitivity to 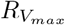(Fig. 3C and Supplementary Fig. S2B, C, D). Very small 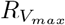, in particular, resulted in a negative mean volume flow rate (Supplementary Fig. S2C) meaning that backflow exceeded forward flow. This suggests that the closure of the valve is crucial for prograde net transport, even more so than the pressure difference required to open the valve or the hydraulic resistance of the open valve (for the parameter ranges considered here, anyway). We also found the mean volume flow rate increases with higher *M*, lower *P*_*d*_, and is insensitive to changes in *D*_*d*_ (Fig. 3D and Supplementary Fig. S2A, E, F). Hence, higher active tension and reduced vessel stiffness increase net transport. This suggests that active properties are more important than passive properties for enhancing transport, and this is achieved by increasing vessel wall compliance with respect to the action of smooth muscle cells. A more detailed analysis of intrinsic pumping is discussed in the next section.

From this analysis, we identified the following parameters that lead to simulation results that align closely with experimental data: 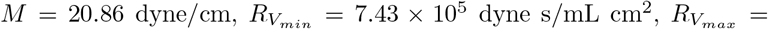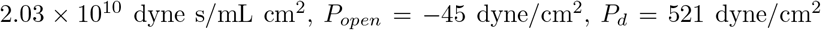. and *D*_*d*_ = 0.015 cm. This parameter set will be used below for further analysis.

### CSF Efflux Through CLVs

The adverse pressure difference that CLVs must overcome is higher than other lymphatic regions. The CSF must be transported against this adverse pressure, and this is enabled through intrinsic pumping in the presence of bileaflet valves in the lymphangions. Fig. 4A-D illustrate this process. In Fig. 4A, the ICP is indicated by a red circle, while central venous blood pressure is represented by a cyan circle. The colors of the X symbols in Fig. 4A correspond to the locations of pressures (Fig. 4B) and diameters (Fig. 4C) for each lymphangion. The arrows in Fig. 4A represent the volume flow rate downstream from the X symbols, and these flow rates are plotted in Fig. 4D using the corresponding colors. It is worth noting that three pressures per lymphangion were calculated, but only the pressures in the middle of each lymphangion were plotted in Fig. 4B to avoid overcrowding. The three pressures computed for each lymphangion are not identical but are similar to each other.

**Fig. 4:**
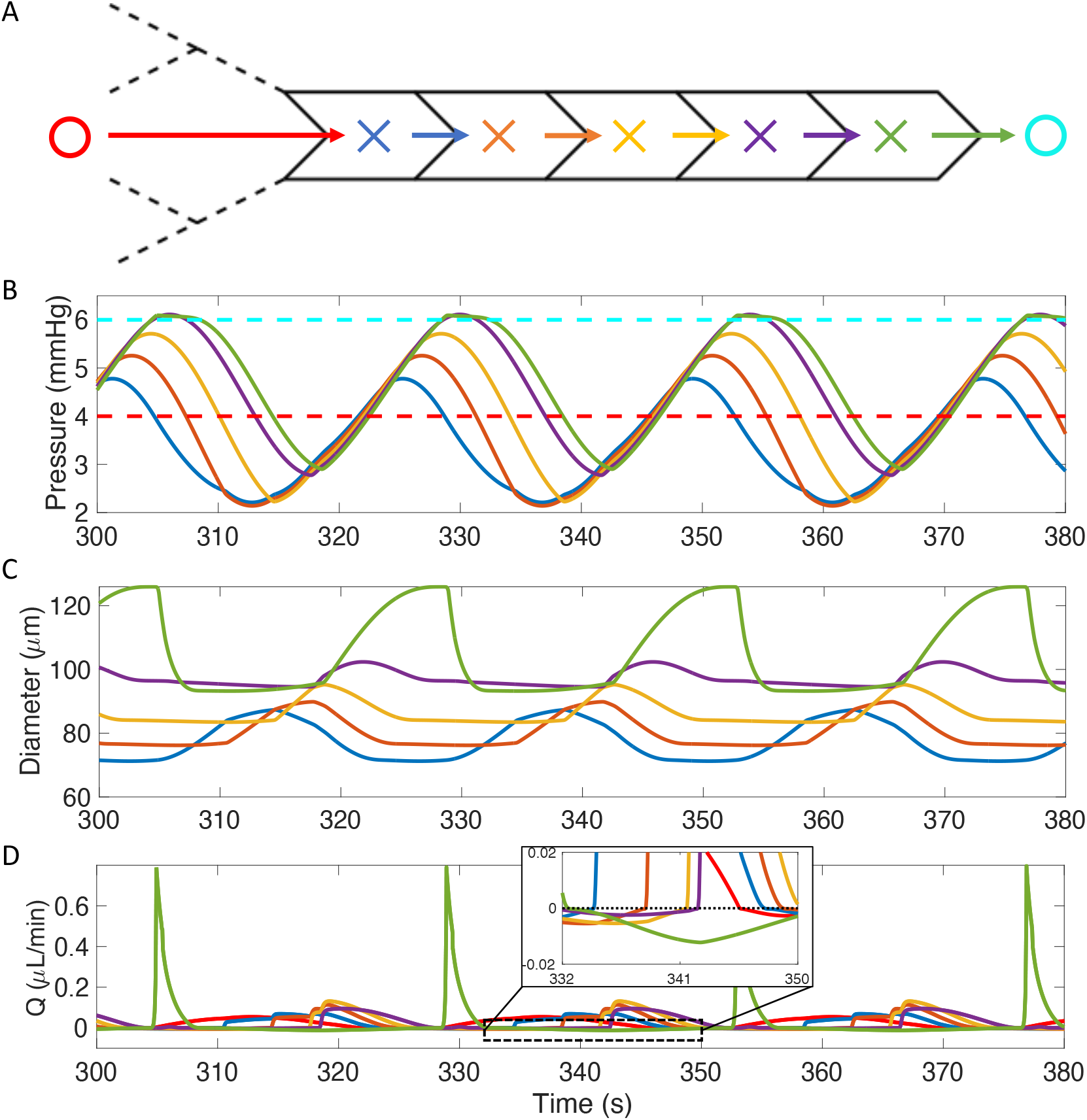
CSF drainage through CLVs. (A) Illustration of the geometry of the simulation. A red and a cyan circle indicate the locations at which upstream pressure (ICP) and downstream pressure (venous blood pressure), respectively, are plotted in (B). Additional X symbols inside each lymphangion are color-coded to indicate the locations of pressure and diameter in (B) and (C); similarly, arrows inside each valve are color-coded to indicate volume flow rate plotted in (D). (B) Plot of pressures in the middle of each lymphangion (*P*_*j,m*_) versus time. We assume constant pressure for the inlet (red dashed line) and outlet (cyan dashed line). (C) Plot of diameters versus time. (D) Plot of volume flow rate versus time. Note the large green peak corresponds to flow through the last valve, furthest right in (A). The subplot in (D) provides a magnified view of the volume flow rate plot for 332 − 350 s to better visualize the backflow.

We now examine the first (blue in Fig. 4A) and second (orange in Fig. 4A) lymphangions. Initially, the pressure in the first lymphangion is lower than the second lymphangion (Fig. 4B). As the pressure inside the first lymphangion begins to exceed that of the second lymphangion (Fig. 4B) and surpasses the additional threshold pressure needed to open the valve (*p*_*open*_), CSF is transported from the first to the second lymphangion (Fig. 4D) as the first lymphangion contracts (Fig. 4C) and opens the valve between them. This results in the expansion of the second lymphangion as the CSF flows in. However, this forward flow is not continuous. The pressure in the second lymphangion exceeds that of the first lymphangion again (Fig. 4B), causing the valve between them close, leading to a small but non-negligible backflow from the second to the first lymphangion (subplot in Fig. 4D). The cycle of expansion and contraction of each lymphangion, driven by oscillatory active tension from smooth muscle cells (intrinsic pumping) and the opening and closing of the valves, enable CSF transport against a relatively large adverse pressure difference (∼2 mmHg), ultimately facilitating fluid drainage into the central venous blood.

An interesting flow is observed at the inlet and outlet of the CLV (Fig. 4D). The volume flow rate at the first valve is positive over a wide time span (red curve in Fig. 4D), due to the presence of large initial lymphatics attached to it. The addition of the net hydraulic resistance of the initial lymphatics results in a gradual change in the volume flow rate. The impact of the net hydraulic resistance of the initial lymphatic vessels will be discussed in more detail in the following section. The outflow from the last lymphangion to the central venous blood is narrow but high (green curve in Fig. 4D). This is due to the higher contraction amplitude of the last lymphangion compared to the other lymphangions. It is important to note that the time-averaged volume flow rate across all lymphangions remains consistent, in accordance with the mass conservation.

### Effect of Intrinsic Pumping and Variable Pressures

Lymphatic vessels rely on both intrinsic and extrinsic pumping mechanisms to transport fluid. Active tension generated by smooth muscle cells in response to stimuli, such as adrenergic and nitric oxide signals, is the primary driver of intrinsic pumping, regulating lymph flow. As active tension (*M*) increases, lymphangion contraction amplitude, which we define as the difference between the maximum and minimum diameter divided by the maximum diameter 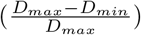, also increases (blue curve in Fig. 5A).

**Fig. 5:**
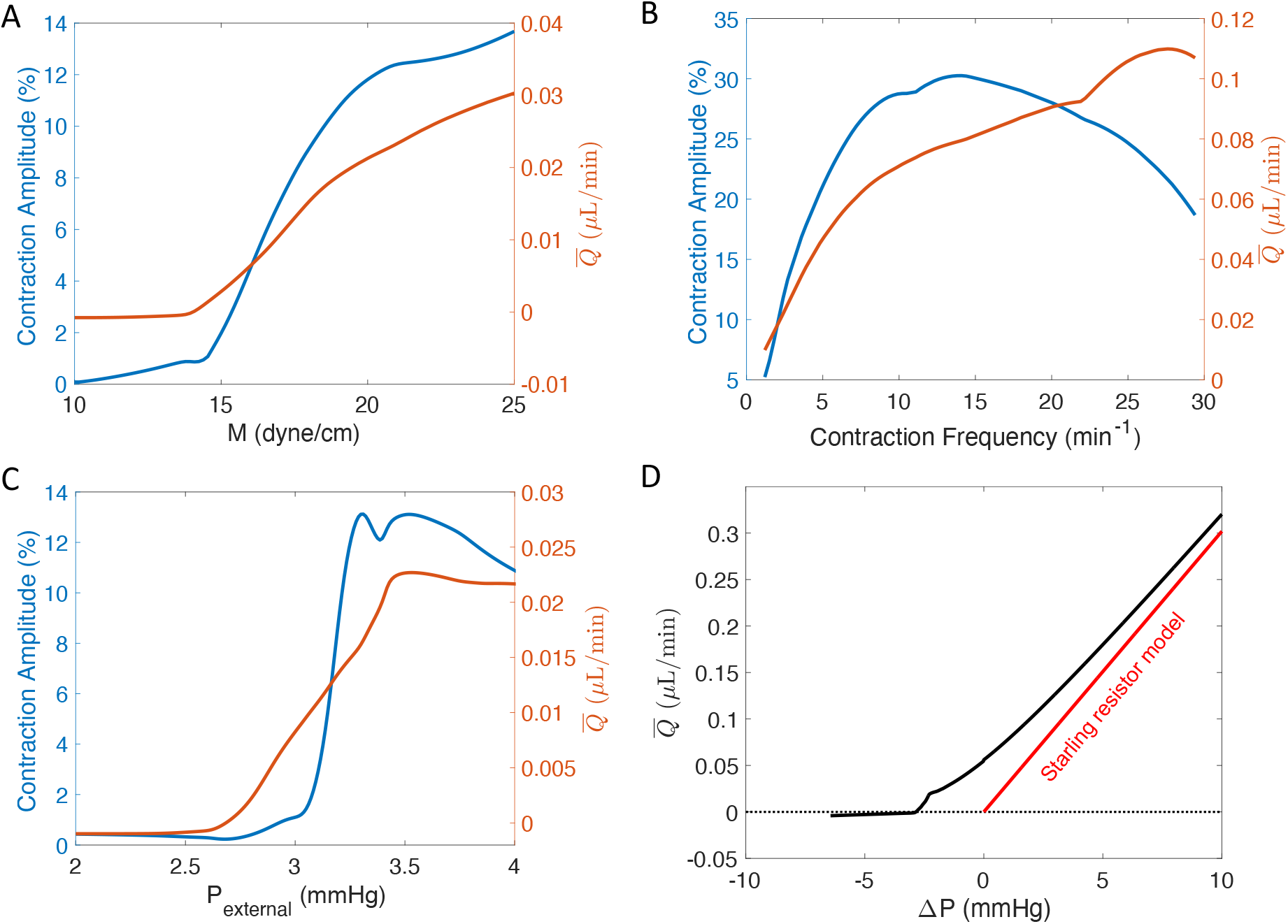
Effects of varying active tension, contraction frequency, external pressure, and longitudinal pressure gradient. These quantities are varied for all lymphangions in our model, but results are plotted for only the third lymphangion. (A) Contraction amplitude (left y-axis) and mean volume flow rate *Q* (right y-axis) as a function of *M* (active tension of smooth muscle cells). (B) Contraction amplitude (left y-axis) and mean volume flow rate 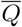(right y-axis) as a function of *f* (contraction frequency). (C) Contraction amplitude (left y-axis) and 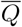(right y-axis) as a function of *P*_*external*_ (external pressure); the longitudinal pressure difference is fixed at Δ*P* = −2 mmHg. (D) 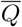 as a function of longitudinal pressure difference Δ*P* = ICP − *P*_*venous*_ (i.e., inlet pressure − outlet pressure); the external pressure is fixed at *P*_*external*_ = 3.75 mmHg. At large Δ*p*, the model volume flow rate approaches that of a Starling resistor (red line), as expected (see main text).

The orange curve in Fig. 5A illustrates the relationship between mean volume flow rate and *M*, indicating that a higher value of *M* (i.e., higher contraction amplitude) results in greater net transport.

In Fig. 5A, two key points stand out. First, as *M* increases in the 10 − 14 dyne/cm range, the contraction amplitude increases slowly, and the mean volume flow rate remains nearly steady. This is because the active tension is too low to significantly affect contractions compared to the lymphangion’s passive properties. In the 15 − 20 range for *M*, the contraction amplitude increases dramatically as active tension dominates the lymphangion’s passive properties. Beyond *M* ≈ 21, the increase in contraction amplitude slows down, suggesting volume flow rate may stabilize beyond this point. This is because the vessel diameter remains small when active tension is large (and hydraulic resistance depends sensitively on diameter, scaling with *D*^4^). This indicates that excessively high tension does not necessary promote CSF transport, and the the optimal range is likely around 20 *dyne/cm*.

Next, we fixed *M* = 20.86 dyne/cm and adjusted the contraction frequency, *f* (Fig. 5B). Initially, the contraction amplitude and the mean volume flow rate rose sharply for frequencies in the approximate range 2 − 14 min^−1^. For frequencies in the range of 14 − 27 min^−1^, the contraction amplitude decreases, but the mean volume flow rate continues to increase. Once the frequency exceeds 27 min^−1^, the volume flow rate begins to decrease slightly. This suggest that there is a wide range over which increases in contraction frequency lead to increases in mean volume flow rate, but transport is most strongly altered for frequencies below about 10 min^−1^. At higher contraction frequencies (≳ 14 min^−1^), the lymphangion’s contraction amplitude is reduced because it begins contracting before becoming fully inflated.

We also tested the effect of varying external pressure (Fig. 5C). When the external pressure is in the lower range (2−2.7) mmHg, the contraction amplitude of the vessels is low, resulting in a low volume flow rate. For external pressures of about 2.7 − 3.3 mmHg, the contraction amplitude of the vessels increases, leading to a corresponding increase in the volume flow rate. However, once the external pressure exceeds about 3.5 mmHg, the contraction amplitude begins to decrease, causing a slight decrease in the volume flow rate. These results indicate that external pressure affects the intrinsic pumping of the CLVs, with an optimal range of external pressure necessary for effective pumping. At lower external pressures, the lymphangion is inflated with reduced contractility and thus the volume flow rate is low. Conversely, at higher external pressures, the diameter of the lymphangion is significantly reduced, but the substantial contraction amplitudes are able to maintain considerable volume flow rate, at least for the range of external pressures we considered.

Finally, we tested different net pressure drops between the inlet and outlet in our model of the CSF outflow pathway (Fig. 5D). We varied both the ICP (inlet pressure) and *P*_*venous*_ (outlet pressure) to model outflow during various scenarios that affect pressure (e.g., different body postures, variable hydration states). From the ranges of ICP and *P*_*venous*_ listed in Table 1), we estimated the pressure drop (Δ*P* = *ICP* − *P*_*venous*_) across the model to vary from −6.44 mmHg to 2.84 mmHg. For Δ*P* values between −6.44 mmHg and −2.9 mmHg, either the inlet pressure was too small or the outlet pressure was too high, resulting in an overall negative mean volume flow rate. This indicates that for this range of pressure differences, the CLV pumping is not strong enough to transport fluid to the cervical lymph nodes and instead a small amount of fluid is transported retrograde. For Δ*P* values above −2.9 mmHg, a positive volume flow rate with a gradually varying slope was observed. Since the external pressure was fixed at *P*_*external*_ = 3.75 mmHg in these simulations, and *P*_*venous*_ ≥ 4.16 mmHg (Table 2), we may conclude that our model should behave like a Starling resistor as Δ*P* increases [50], i.e., the volume flow rate 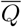 should vary in proportion with Δ*P*. We analytically computed the volume flow rate as a function of Δ*P* for a Hagen-Poiseuille flow through a tube (with length 5*L* and diameter matching the temporal mean diameter of the third lymphangion for each given Δ*P* value) connected in series with a hydraulic resistance *R*_*initial*_; the result is plotted in red in Fig. 5D. As expected, our model approaches that of the Starling resistor as Δ*P* increases. This provides both a source of model verification and insight that at large favorable pressure gradients, the external pressure becomes unimportant.

### Effect of Lymphatic Capillary Branching Assumptions

In our final test, we examined how lymphatic capillary branching affects net volume flow rate. We applied an inlet pressure corresponding to either normal ICP (4 mmHg) or elevated ICP (10 mmHg), as indicated in Fig. 6A. We then tested two different branching scenarios: one adhering to the typical Murray’s law (with exponent 3) and another corresponding to Murray’s law with exponent 1.45 [39]. The model with the smaller exponent led to smaller vessel diameters, but longer length, across each bifurcation compared to the normal case, ultimately generating a higher hydraulic resistance *R*_*initial*_.

**Fig. 6:**
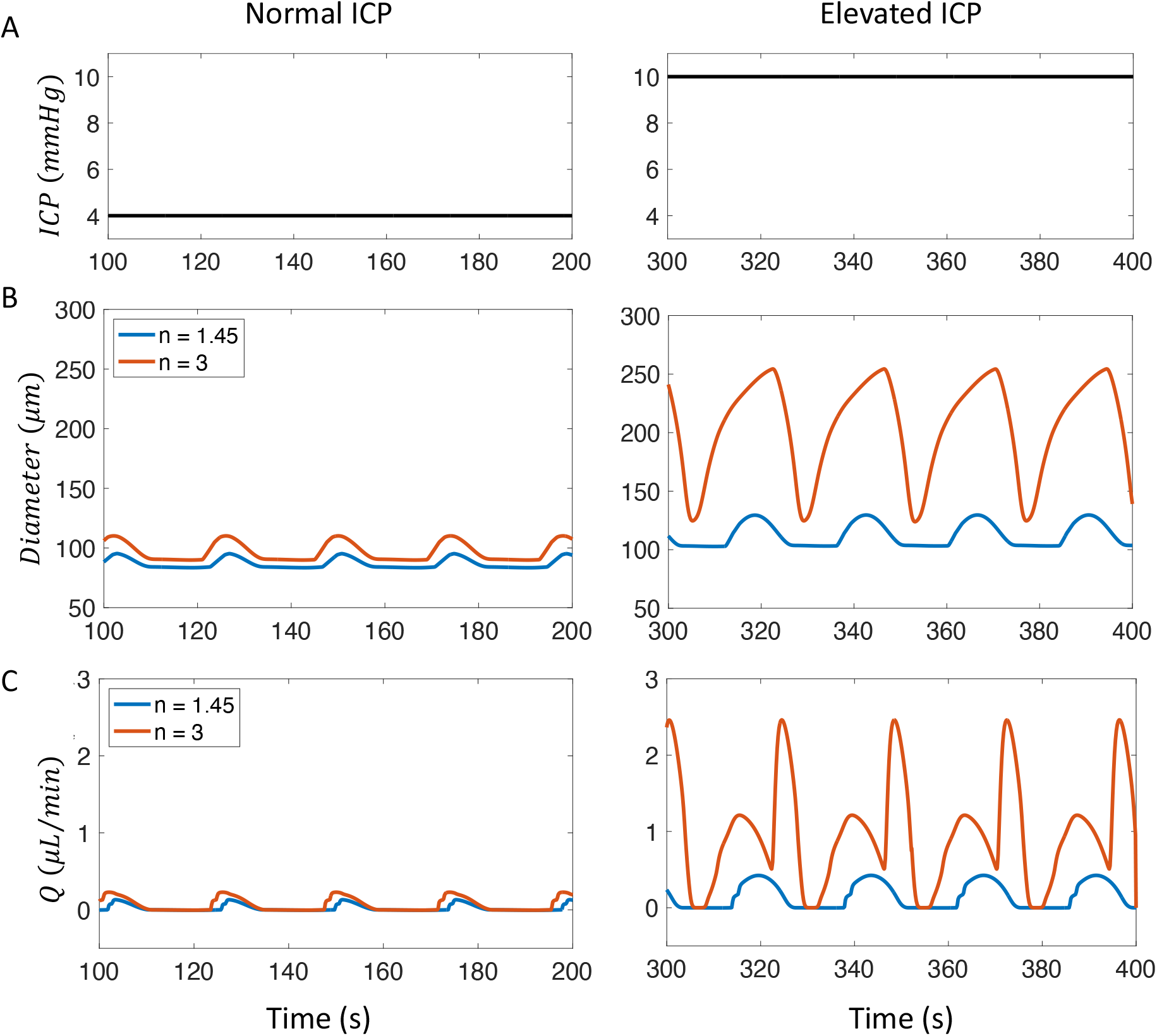
Impact of lymphatic capillary branching on diameter and volume flow rate under different ICP conditions. (A) Plots of inlet pressure (ICP) for a normal (left) and elevated (right) case. (B) CLV diameter versus time for different ICP. The exponent in Murray’s law is *n* = 1.45 (blue curve) or *n* = 3 (orange curve). (C) Volume flow rate versus time for different ICP. Note that the value *n* = 1.45 comes from an experimental study of murine dermal lymphatic capillaries [39].

At normal ICP, both the diameter and volume flow rate for the two cases of different exponents operate within a physiological range. The diameter and volume flow rate for typical Murray’s law are only slightly higher than those for the modified Murray’s law (Fig.6B,C). However, at elevated ICP, the differences for the two different exponent cases become substantial. Both the diameter and volume flow rate increase for both exponent cases (Fig.6B and C), but the model with modified Murray’s law (*n* = 1.45, blue curve) shows only a small increase in diameter which spans a range of about 100 to 130 µm. On the other hand, the model with normal Murray’s law (*n* = 3, orange curve) exhibits large amplitude fluctuations, ranging from about 130 to 250 µm. Physiologically, such a large peak diameter and range in diameter is unrealistic and could likely lead to vessel rupture when ICP increases. The volume flow rate also increases with the elevated ICP, as expected when the upstream pressure exceeds the downstream pressure. However, the model with the modified Murray’s law (*n* = 1.45, blue curve) shows a gradual increase in volume flow rate, while the model with normal Murray’s law (*n* = 3, orange curve) exhibits an extreme increase. Excessive outflow due to elevated ICP would likely have deleterious effects. These results suggest that lymphatic capillaries that absorb the CSF in the SAS likely do not follow normal Murray’s law but may instead adhere to a modified version with a smaller exponent (*n <* 3) so that the associated increase in hydraulic resistance will buffer transport rates with the respect to changes in ICP.

## Discussion

Lymphatic vessels transport fluid against an adverse pressure gradient, and CLVs are no exception. In fact, CLVs must overcome a higher adverse pressure gradient compared to other lymphatics [46, 47], which is a consequence of their anatomy which bridges the skull and cervical lymph nodes. Our numerical model suggests that both active and passive properties of the vessel wall and the hydraulic resistance of the valve in its closed state are crucial for maintaining the vessel’s diameter and volume flow rate. In particular, we found that transport is sensitive to maximum valve resistance 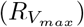 when that value is small (Fig. 3C). This sensitivity results because, if the valves do not close completely (i.e., 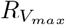 is relatively small), substantial backflow will occur leading to a diminished net volume flow rate. Recent *in vivo* measurements of CLV flow in mice [12, 51] support the conclusions drawn here regarding the most sensitive parameters. Our simulations predict volume flow rates through a single CLV that range from about 0 to 0.1 *µ*L/min as active tension or contraction frequency are varied (Fig. 5A-B), which is in good agreement with CLV volume flow rates reported by Du et al. [51]. They found that volume flow rates decrease from 0.071 to 0.017 *µ*L/min for 2-month-old versus 22-month-old mice [51]. Furthermore, they attributed this reduced transport primarily to a decrease in contraction frequency, as well as valve dysfunction that increased retrograde flow, reducing net transport.

As a lymphangion expands, the pressure inside it decreases, allowing cerebrospinal fluid (CSF) to enter. As it contracts, the CSF is transported to the next lymphangion, opening the valve in between and eventually draining to the central venous blood. It is noteworthy that the average diameters of each of the five lymphangions we model increase as one moves in the downstream direction. This is because the same parameters were used for all five lymphangions and the transmural pressure difference increases along the streamwise direction. Parameters that control the passive properties of the wall, such as distensibility (*D*_*d*_) and the pressure at which the vessel is fully distended (*P*_*d*_), can be modeled to increase along the downstream direction [28, 40] in order to maintain the same average diameters. Our results highlight differences between CLVs and mesenteric lymphatic vessels, in that CLVs exhibit a stronger sensitivity to wall stiffening (*D*_*d*_ and *P*_*d*_) with changes in diameter at both positive and negative transmural pressures [28]. Due to this stiffer wall, the smooth muscle cells need to generate stronger active tension (*M*) to transport CSF.

In the study by Hussain et al. it was observed that the CSF outflow through CLVs decreases following TBI [12]. This study also found that the level of norepinephrine increased after TBI, which was associated with a decrease in the contraction amplitude and alteration to the frequency of the CLVs. Our simulations support the idea that decreased contraction amplitude and/or frequency lead to a decreased volume flow rate. This suggests that enhancing lymphatic activity may provide an effective target for restoring CSF drainage and enhancing brain waste clearance, as recently demonstrated by Du et al. [51]. Currently, our model only includes intrinsic pumping, which captures net volume flow rates that are in good agreement with experimental measurements. However, future numerical and/or experimental work may investigate the role of external pressures, such as those exerted by skeletal muscles or arising from neck massage, which may enhance drainage.

Various external forces act on the CLVs from sources such as pulsatility of nearby arteries, movement due to respiration, and skeletal muscle contractions; however, a recent *in vivo* study suggests cardiac pulsatility and respiration does not play an important role in CLV transport [51]. The pressure gradient formed by the difference between the external pressure and the intraluminal pressure affects the contractility of the CLVs. A small external pressure leads to inflation of the vessel with small amplitude contractions, while a large external pressure decreases the vessel diameter but increases the contraction amplitude. This indicates that an optimal range of external pressure likely exists, which is necessary for effective pumping and net transport. Currently, we have tested external pressure in the range of 2 mmHg to 4 mmHg. We expect that beyond about 4 mmHg, total collapse of the vessel is likely because the transmural pressure gradient will become negative. This negative pressure gradient leads to deformation of the vessel wall. However, when we tested pressures beyond 4 mmHg, numerical instability and failure occurred due to the calculation of negative diameter values. It may be helpful to test the effect of external pressure using a different model of the transmural pressure equation 6 in the future.

The ICP, which was assumed as the inlet pressure in our model, can increase due to gross swelling or intracranial bleeding after TBI [52]. One may hypothesize that this increased ICP could lead to rapid drainage of CSF from the skull. However, our simulations indicate that if the initial lymphatics that absorb the CSF bifurcate according to a modified version of Murray’s law (with exponent substantially less than the typical *n* = 3), the associated increase in hydraulic resistance will act as a buffer that inhibits rapid CSF drainage. Murray’s law, with an exponent of 3, was formulated to describe the bifurcations of arteries, aiming to explain how arterial branching minimizes hydraulic resistance. Our simulations suggest that minimizing hydraulic resistance in the initial lymphatic branches connecting the skull and CLVs is perhaps deleterious. Additionally, it is worth noting that the assumptions associated with Murray’s law are already violated due to the presence of the valves, suggesting one should not expect to observe an exponent of *n* = 3 associated with the branching. Future experiments could test for evidence of this safety mechanism by directly measuring the diameters of sequential initial lymphatic vessels or even by counting the number of branching generations. Our calculations suggest that approximately 11 versus 5 generations exist for Murray’s law exponents of *n* = 3 versus *n* = 1.45. We also highlight that the exponent value *n* = 1.45 comes from a study of murine dermal lymphatic capillaries [39], and the true value for nasal lymphatics may differ substantially.

Substantial evidence indicates that CSF drains through the cribriform plate to nasal lymphatics in rodents [13]. However, this pathway appears less important in humans [53], highlighting critical differences that may exist in CSF pathways for rodents versus humans. Meningeal lymphatics may potentially serve as a primary efflux route in humans. In our model, the initial lymphatic vessels are currently based on the nasopharyngeal lymphatic vessels of rodents. However, our model can be adapted to capture efflux to any other lymphatic vessels with different dimensions and properties, including meningeal lymphatic vessels.

In the recent study by Yoon et al., a complex network of lymphatics draining CSF through the nasopharynx to the cervical lymph nodes was identified [54]. This plexus contains tiny lymphatic vasculature at upstream locations, which merge into larger vessels. These larger vessels are considered pre-collector lymphatic vessels due to the presence of valves and somewhat loose covering of smooth muscle cells. Currently, our model does not include pre-collector lymphatic vessels. Future studies could incorporate a model of the nasopharyngeal plexus and test its hydraulic resistance, which may further buffer changes in ICP. Additionally, the presence of valves in the plexus supports the idea that hydraulic resistance of the valve in its closed state is critical for CLVs when the pressure difference is high. Although the closure of lymphatic valves is not perfect and does allows a small amount of backflow, the series of valves located in the plexus likely help reduce backflow, allowing the CLVs to maintain an optimal diameter and volume flow rate.

Several important limitations of this study should be acknowledged. Spatial resolution is limited in our simulation which prevents, for example, detailed simulation of variation in contraction amplitude along the length of each lymphangion. As a consequence of using lumped parameter modeling, our predictions only include volume flow rates and pressures for three points per lymphangion (one at the center, and one very close to each valve). However, a more detailed treatment is precluded by the current scarcity of quantitative data characterizing CLV geometry and physical properties. A further limitation is that the lumped parameter approach assumes that contractions of adjacent lymphangions occur with some fixed temporal phase (parameter *t*_*d*_ in Table 1). A more detailed analysis of relation between flow and contraction timings, including both the frequency and *t*_*d*_, will be conducted in future work. *In vitro* observations of CLV contractions following exposure to norepinephine reveal loss of contraction entrainment [12], which cannot be readily modeled using the lumped parameter approach. Nonetheless, the lumped parameter method is appealing in that simulations are fast and our Monte Carlo approach could be conducted efficiently. Overall, our study has uncovered valuable insights into not only fundamentals of the physical properties of CLVs, but also potential therapeutic strategies.

## Conclusions

In this study, we performed simulations of CSF efflux to cervical lymphatic vessels using a lumped parameter model. This model captures the CSF drainage from the subarachnoid space, absorbed by the initial lymphatics embedded in the nasal region, which merge over several generations and drain to the cervical lymphatic vessels, eventually reaching the central venous blood. We identified parameters that are unknown and difficult to experimentally measure using a Monte Carlo search in which we matched simulation predictions to *in vivo* measurements. Simultaneously, we explored how the unknown parameters in the governing equations affect the median diameter of the vessels and the mean volume flow rate, concluding that magnitude of active tension, passive properties of the vessel wall, and hydraulic resistance of the valve in its closed state have the greatest effect. We also demonstrated that increasing the active tension and/or the contraction frequency increases the overall volume flow rate, and we tested how the bifurcations of the upstream lymphatic capillaries affect the overall flow in response to elevated ICP. Narrower and longer branches (arising from a modified form of Murray’s law with exponent 1.45 [39]) increases the net hydraulic resistance, increasing the robustness of the system to elevated ICP.

This is the first numerical study of CSF drainage through CLVs. We anticipate that our rigorous parameter analysis using our proposed model will help guide future numerical studies aimed at modeling CLVs under physiological and pathological conditions. Additionally, our results form a foundation for future experiments in this field, contributing to our understanding of CSF drainage and its potential therapeutic applications.

## Appendix A Derivation of Lumped Parameter Equations

In this appendix, we present the derivation of lumped parameter fluid flow equations that are implemented as our governing equations. The equations for transmural pressure (equation 6) and valve resistance (equation 4) are directly adopted from the previous works of Bertram el al. [28, 40]. Therefore, the derivations of those equations are not included here. The mass conservation for 1D flow can be written as:

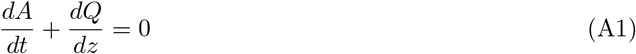

where *A*(*z, t*) is the cross-sectional area of the vessel and *Q*(*z, t*) is the volume flow rate. Equation (A1) can be integrated over one lymphangion of length *z*_2_ − *z*_1_ = *L*:

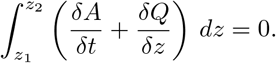

This integration can be split into two parts:

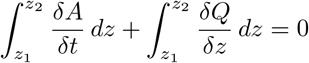

The first part can be computed as:

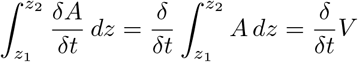

where *V* (*t*) is the lymphangion volume. The second part can be computed as:

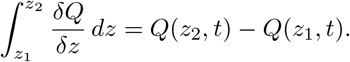

If we denote *Q*_*j*+1_ = *Q*(*z*_2_, *t*) and *Q*_*j*_ = *Q*(*z*_1_, *t*), where *j* indexes the valves separating each lymphangion, then the equation can be rewritten in the form:

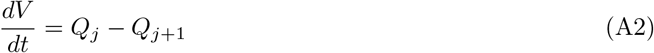

If we assume the lymphangion has a circular cross-sectional area over a length *L*, then the volume *V* can be written as 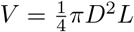. The temporal derivative of *V* is then 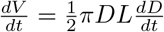. Thus, the lumped parameter mass conservation equation (A2) can be written in the form given by equation (2) in the main text:

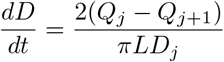

The conservation of momentum equation can be written as follows, with the assumptions that the flow is quasi-steady and laminar, the velocity in the radial direction is zero (*v*_*r*_ = 0), the axial velocity is independent of the circumferential direction 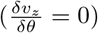, and the pressure gradient is constant over the length of a given lymphangion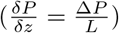:

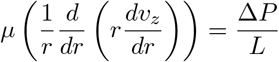

After integrating with respect to *r* and applying the boundary conditions (*v*_*z*_ is finite at *r* = 0 and *v*_*z*_ = 0 at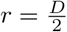), the equation can be written as:

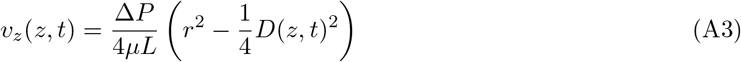

where *µ* is the dynamic viscosity of the fluid. The volume flow rate, *Q*, is the integral of the axial velocity over the cross-sectional area of the pipe:

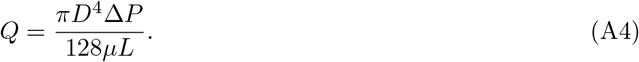

After integration, we obtain the Hagen-Poiseuille equation:

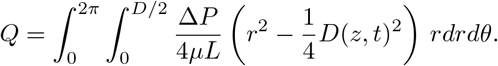

Since the midpoint pressure inside a lymphangion (*P*_*j,m*_) is used to calculate the pressure drop, *L/*2 should be used instead of *L*. Thus, equation (3) in the main text is obtained:

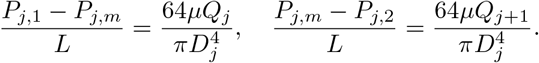

## Ethics approval and consent to participate

Not applicable.

## Consent for publication

Not applicable.

## Availability of data and materials

Data and materials are available from the authors following any reasonable request. Simulation codes are included as supplementary material.

## Authors’ contributions

Conceptualization: DK, JT; Methodology: DK, JT; Formal analysis and investigation: DK; Writing - original draft preparation: DK; Writing - review and editing: JT; Funding acquisition: JT; Resources: JT; Supervision: JT

## Acknowledgements

Not applicable.

## Supplementary information

**Fig. S1:**
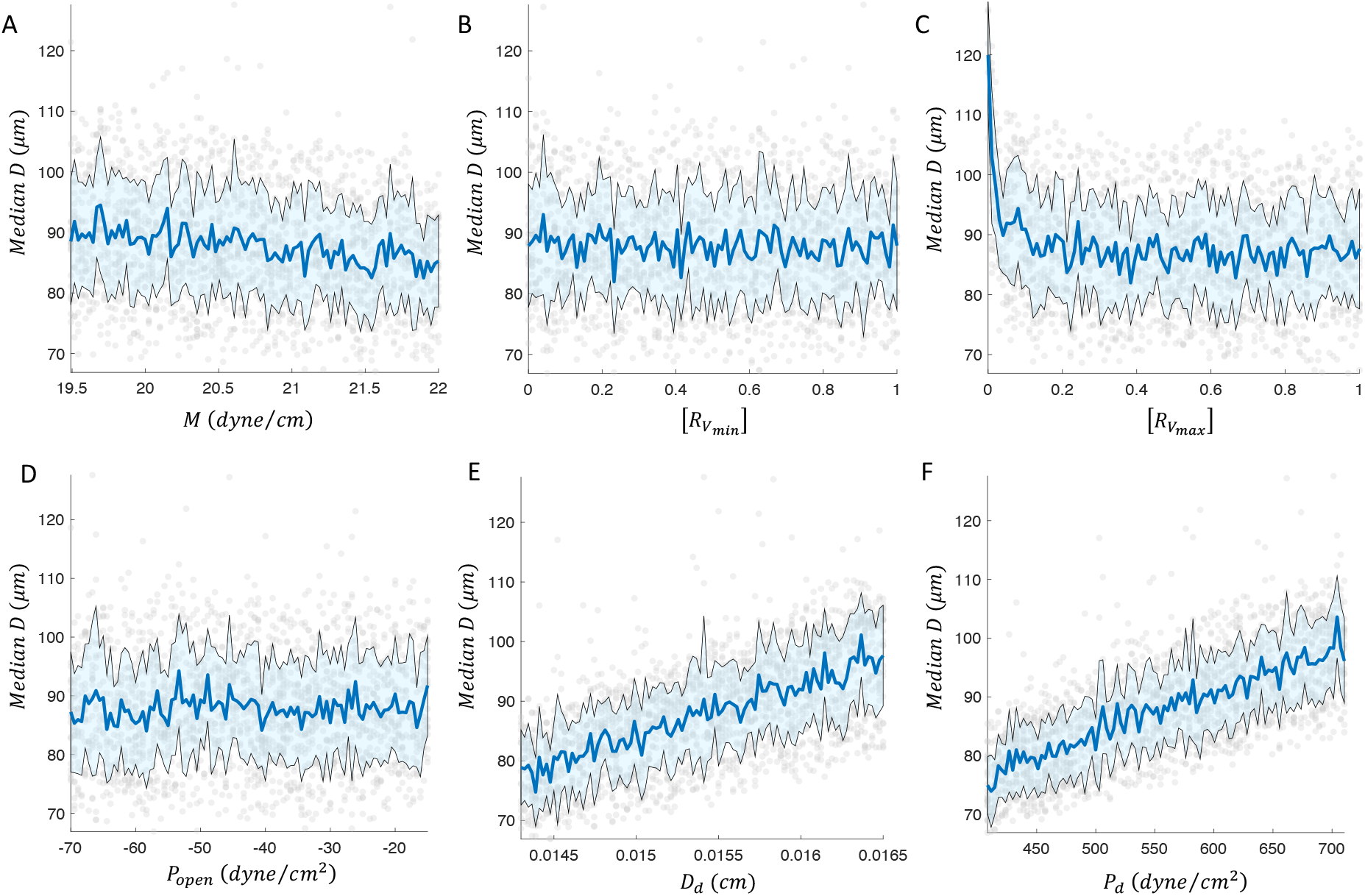
Supplementary analysis of parameter sensitivity in median diameter. All data points are represented as light grey scattered points. The mean value is plotted in blue, while mean plus/minus one standard deviation is plotted in black (and the region in between is shaded light blue). (A-F) Median diameter as a function of (A) active tension, (B) normalized hydraulic resistance of the open valve, (C) normalized hydraulic resistance of the closed valve, (D) threshold pressure to open the valve, (E) threshold diameter for positive or negative transmural pressure in the vessel, and (F) vessel wall stiffness.

**Fig. S2:**
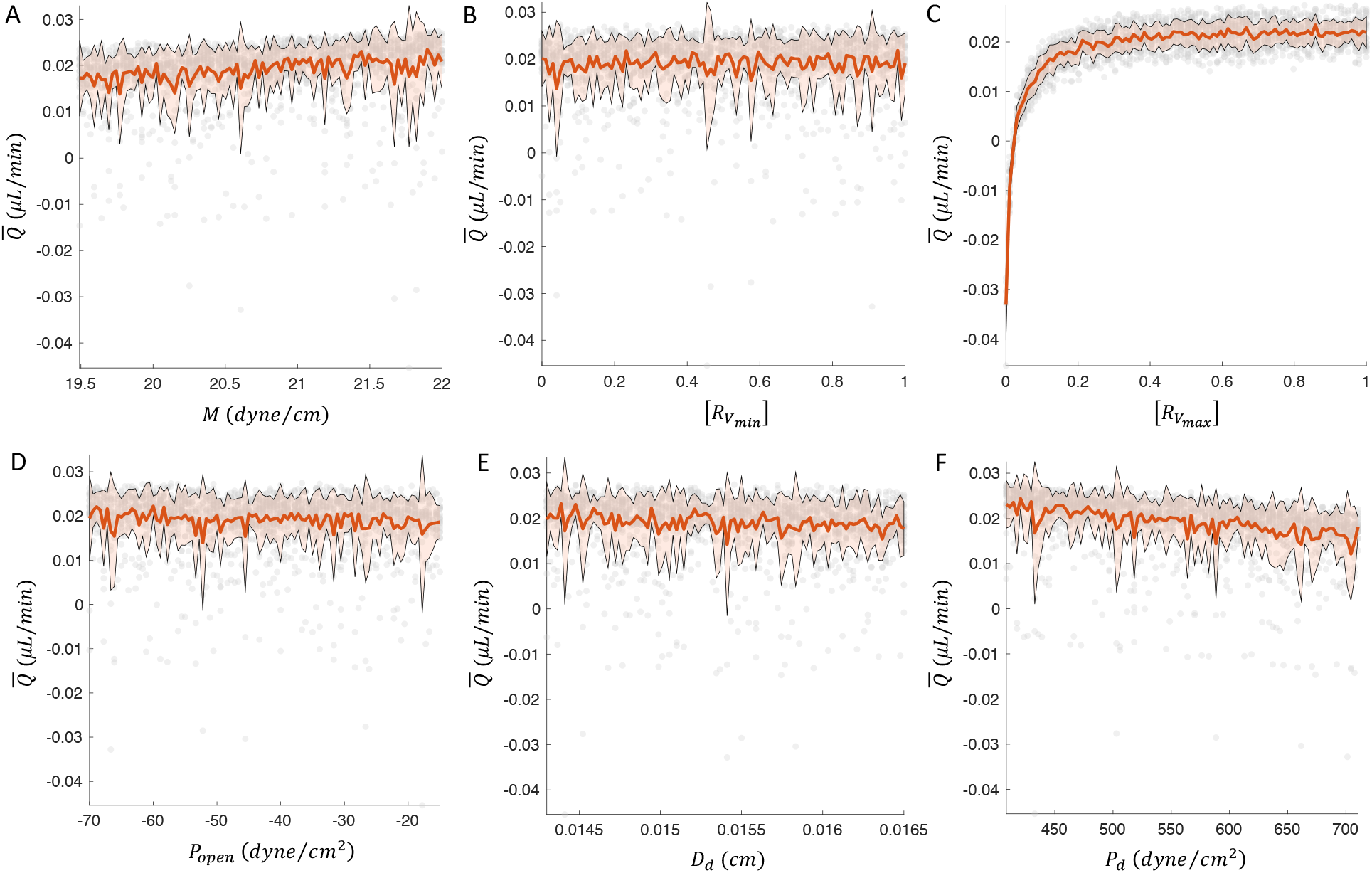
Supplementary analysis of parameter sensitivity in mean volume flow rate. All data points are represented as light grey scatter points. The mean value is plotted in orange, while mean plus/minus one standard deviation is plotted in black (and the region in between is shaded light orange). (A-F) Mean volume flow rate as a function of (A) active tension, (B) normalized hydraulic resistance of the open valve, (C) normalized hydraulic resistance of the closed valve, (D) threshold pressure to open the valve, (E) threshold diameter for positive or negative transmural pressure in the vessel, and (F) vessel wall stiffness.

